# Effects of Repeated Treatment with Monoamine-Transporter-Inhibitor Antidepressants on Pain-Related Depression of Intracranial Self-Stimulation in Rats

**DOI:** 10.1101/753905

**Authors:** LP Legakis, L Karim-Nejad, SS Negus

**Author notes:** Communicating Author: S. Stevens Negus.

## Abstract

Synaptic neurotransmission with dopamine (DA), norepinephrine (NE), and serotonin (5-HT) is terminated primarily by reuptake into the presynaptic terminal via the DA, NE, and 5-HT transporters (DAT/NET/SERT, respectively). Monoamine transporter inhibitors constitute one class of drugs used to treat pain, and emergence of analgesic effects by these compounds often requires repeated treatment for days or weeks. The present study compared antinociceptive effects produced by repeated treatment with monoamine transporter inhibitors in a preclinical assay of pain-related depression of positively reinforced operant responding. Adult male Sprague-Dawley rats equipped with microelectrodes targeting a brain-reward area responded for pulses of electrical brain stimulation in an intracranial self-stimulation (ICSS) procedure. Intraperitoneal injection of dilute lactic acid served as a noxious stimulus that repeatedly depressed ICSS and also produced weight loss during 7 days of repeated acid administration. Both acid-induced ICSS depression and weight loss were blocked by repeated pretreatment with the nonsteroidal anti-inflammatory drug ketorolac (a positive control) but not by the kappa opioid receptor agonist U69,593 (a negative control). Like ketorolac, the DAT/NET inhibitor bupropion fully blocked acid-induced ICSS depression and weight loss throughout all 7 days of treatment. Conversely, the NET-selective inhibitor nortriptyline and SERT-selective inhibitor citalopram produced antinociception only after several days of repeated treatment, and weight loss was attenuated by citalopram but not by nortriptyline. These results support effectiveness of bupropion to alleviate signs of pain-related behavioral depression in rats and further suggest that nortriptyline and citalopram produce a more gradual onset of antinociception during repeated treatment.

## INTRODUCTION

Mu opioid receptor agonists (e.g. morphine) and nonsteroidal anti-inflammatory drugs (NSAIDs, e.g. ketorolac) are among the most widely used analgesics for treatment of moderate to severe pain, but they are not always effective, and their use is often constrained by side effects (Litvak and McEvoy 1990; Matava 2018; Yaksh and Wallace 2018). Drugs that inhibit the norepinephrine transporter (NET), serotonin transporter (SERT), and/or dopamine transporter (DAT) represent another class of drugs that is sometimes used to treat pain (Obata 2017; Sutherland et al. 2018). Norepinephrine (NE), serotonin (5-HT), and dopamine (DA) are monoamine neurotransmitters involved in a wide range of physiological and behavioral processes (Jacob and Nienborg 2018; Nutt 2008). Monoamine transporters located on presynaptic terminals are the primary mechanism for neurotransmitter clearance from a synapse after monoamine release, and transporter inhibition reduces neurotransmitter clearance, increases synaptic neurotransmitter concentrations, and increases signaling via the associated monoamine receptors (Aggarwal and Mortensen 2017; Lin et al. 2011). Monoamine transporter inhibitors are most widely used for the treatment of major depression (Cipriani et al. 2018; O’Donnell et al. 2018); however, pain is often associated with depression-like signs and symptoms, and at least some dimensions of pain may be mediated by changes in monoamine signaling similar to those that are also present in major depression (Boakye et al. 2016; Goesling et al. 2013). The effectiveness of monoamine transporter inhibitors for pain treatment was first established with so-called “tricyclic” antidepressants, and tricyclics such amitriptyline and its primary metabolite nortriptyline, which act primarily as NET inhibitors, continue to be used (Finnerup et al. 2015; Moore et al. 2015; Paoli et al. 1960). However, these compounds have low selectivity for monoamine transporters vs. non-transporter targets such as cholinergic receptors, and some of their side effects are mediated by these non-transporter targets (O’Donnell et al. 2018). Several more recently developed drugs display greater selectivity for monoamine transporters vs. non-transporter targets and may act either simultaneously at multiple transporters (e.g. the NET/SERT inhibitor duloxetine or DAT/NET inhibitor bupropion) or selectively at a single transporter (e.g. the SERT-selective inhibitor citalopram) (Bymaster et al. 2005; Hyttel et al. 1992; O’Donnell et al. 2018; Stahl et al. 2004). Analgesic effectiveness is best established for NET/SERT inhibitors (Attal 2019; Wang et al. 2015), but DAT/NET inhibitors (Pud et al. 2017; Shah and Moradimehr 2010) and SERT-selective inhibitors (Barakat et al. 2018; Lunn et al. 2015) may also be effective under at least some conditions.

Monoamine transporter inhibitors have been reported previously to produce antinociception in preclinical laboratory-animal procedures that rely on “pain-stimulated behaviors,” which can be defined as behaviors that increase in rate, frequency, or intensity after delivery of a putative pain stimulus (e.g. paw or tail withdrawal from thermal or mechanical stimuli) (Gatch et al. 1998; Hall et al. 2011; Pedersen et al. 2005; Ventafridda et al. 1990). However, pain states can also be associated with decreases in behavior, and pain-related behavioral depression is both a common criterion of pain diagnosis and target of pain treatment in both human and veterinary medicine (Brown et al. 2008; Dworkin et al. 2005). Accordingly, we and others have developed preclinical assays of pain-depressed behaviors, which can be defined as behaviors that decrease in rate, frequency, or intensity after delivery of a pain stimulus (Negus 2019; Tappe-Theodor et al. 2019). As one example, we reported previously that intraperitoneal delivery of dilute lactic acid (IP acid) could serve as a noxious stimulus to decrease positively reinforced operant responding maintained by delivery of rewarding electrical brain stimulation in an intracranial self-stimulation (ICSS) procedure (Negus 2013). IP acid-induced depression of ICSS can be alleviated by acute treatment with clinically effective opioid and NSAID analgesics, but not by clinically ineffective classes of analgesics (e.g. centrally active kappa opioid receptor agonists such as U69,593) (Altarifi et al. 2015; Negus et al. 2010; Pereira Do Carmo et al. 2009). Among monoamine transporter inhibitors, acute treatment with bupropion was also effective to relieve IP acid-induced ICSS depression, but NET- and SERT-selective inhibitors (including citalopram) were not (Rosenberg et al. 2013).

A distinguishing characteristic of monoamine transporter inhibitors for pain treatment is that they typically require repeated treatment for days to weeks before analgesic effects emerge (Sutherland et al. 2018). Accordingly, the main goal of the present study was to build on our previous work by evaluating the effectiveness of repeated daily treatment with bupropion, nortriptyline, and citalopram to alleviate IP acid-induced ICSS depression. The effects of these monoamine transporter were compared to effects of ketorolac and U69,593 as positive and negative controls, respectively. All drugs were evaluated in a procedure in which rats received daily treatment with IP acid for 7 days, and we have shown previously that repeated IP acid produces repeatable ICSS depression that can be blocked by daily co-administration of morphine (Miller et al. 2015). We hypothesized that all three monoamine transporter inhibitors would alleviate IP acid-induced ICSS depression after repeated treatment.

## METHODS

### Subjects

Adult male and female Sprague-Dawley rats (Envigo, Somerset, NJ) with initial weights of 376 – 508 g (males) and 266 – 328 g (females) were housed individually and maintained on a 12-h light/dark cycle with lights on from 6:00AM to 6:00PM in an AAALAC International-accredited housing facility. Food and water were available ad libitum in the home cage. Animal-use protocols were approved by the Virginia Commonwealth University Institutional Animal Care and Use Committee and were in accordance with the National Academy of Science’s Guide for the Care and Use of Laboratory Animals (National_Research_Council 2003).

### Intracranial self-stimulation (ICSS)

#### Surgery

Male and female rats (N=20 of each sex) were anesthetized with isoflurane (2.5-3% in oxygen; Webster Veterinary, Phoenix, Arizona, USA) and implanted with electrodes (Plastics One, Roanoke, Virginia, USA) in the left medial forebrain bundle at the level of the lateral hypothalamus using previously published procedures and coordinates (Males: 2.8 mm posterior to bregma, 1.7 mm lateral to the midsagittal suture, 8.8 mm below skull surface; Females: 3.8 mm posterior to bregma, 1.6 mm lateral to the midsaggital suture, 8.7 mm below skull surface (Miller et al. 2015; Negus and Miller 2014). The electrode was secured to the skull with orthodontic resin and skull screws. Ketoprofen (Spectrum Chemical, New Brunswick, NJ, 5 mg/kg) was administered immediately and 24 hours after surgery as a postoperative analgesic, and rats recovered for 7 days prior to initiation of ICSS training.

#### Apparatus

Studies were conducted in sound-attenuating boxes containing modular acrylic and metal test chambers (29.2 × 30.5 × 24.1 cm; Med Associates, St Albans, VT, USA). Each chamber contained a response lever (4.5 cm wide, 2.0 cm deep, 3.0 cm above the floor), three stimulus lights (red, yellow, and green) centered 7.6 cm above the lever, a 2-W house light, and an ICSS stimulator. Electrodes were connected to the stimulator via bipolar cables routed through a swivel commutator (Model SL2C, Plastics One, Roanoke, VA, USA). Computers and interface equipment operated by custom software controlled all operant sessions and data collection (Med Associates).

#### Training

Rats were trained to respond for brain stimulation using procedures identical to those previously described (Miller et al. 2015; Negus and Miller 2014). Briefly, a white house light was illuminated during behavioral sessions, and responding under a fixed-ratio (FR) 1 schedule produced a 500-ms train of 0.1-ms square-wave cathodal pulses together with 500-ms illumination of stimulus lights over the response lever. The terminal schedule consisted of sequential 10-min components. Each component consisted of 10 1-min trials, and the available brain-stimulation frequency decreased in 0.05 log Hz increments from one trial to the next (158-56 Hz). Each frequency trial consisted of a 10-s timeout, during which five noncontingent stimulations were delivered at the frequency available during that trial, followed by a 50-s “response” period, during which responding resulted in electrical stimulation. Training continued with presentation of three sequential components per day until the following two criteria for stable responding were met for three consecutive days: (1) ≤5% variability in the maximum rate of reinforcement in any trial, and (2) ≤10% variability in the total number of stimulations per component.

#### Testing

Experiments were conducted using an 8-day protocol. On Day 0, a three-component ICSS session was conducted to establish pre-drug baseline parameters for the number of total stimulations per component and the maximal control rate (MCR) (see Data Analysis). On Days 1 – 7, rats were weighed, and daily experimental sessions consisted of three daily baseline ICSS components followed first by a timeout period when treatments were administered and then by two ICSS test components. Twelve groups of rats were used to evaluate effects of 12 different treatments. Specifically, six groups received daily injections of vehicle or one of five test drugs followed by treatment with 1.8% lactic acid. The other six groups received the same daily injections of vehicle or test drug followed by treatment with acid vehicle. The test drugs and doses were as follows: 3.2 mg/kg bupropion, 3.2 mg/kg nortriptyline, 10 mg/kg citalopram, 10 mg/kg ketorolac, and 0.18 mg/kg U69,593. The interval between administration of test drug or drug vehicle and subsequent administration of IP acid or acid vehicle was 30 min for vehicle, bupropion, nortriptyline, and citalopram, 15 min for ketorolac, and 10 min for U69-593. These doses and pretreatment times were based on previously published studies that examined acute effects of these drugs on ICSS performance in the absence or presence of the IP acid noxious stimulus (Leitl et al. 2014; Moerke et al. 2019; Rosenberg et al. 2013). The treatment drug or drug vehicle was administered at the beginning of the timeout period during daily ICSS sessions, and IP acid or acid vehicle was administered at the end of the timeout period, immediately before initiation of the ICSS test components.

Each treatment was tested in 6-7 rats. Most rats (35) were included in two different groups, receiving no more than one of the three monoamine uptake inhibitors (bupropion, nortriptyline, or citalopram) and no more than one of the three control treatments (vehicle, ketorolac, or U69,593). Rats tested twice received repeated IP acid with only one treatment and repeated IP acid vehicle with the other treatment. Evaluation of sex differences in treatment effects was not a goal of the present study, but to comply with National Institute of Health guidance for inclusion of both sexes in preclinical research, each group consisted of 3-4 male rats and 3-4 female rats.

#### Data analysis

ICSS data were analyzed as previously described (Miller et al. 2015; Negus and Miller 2014). The primary dependent measure was the total number of reinforcements per component (i.e. the total number of stimulations delivered across all brain-stimulation frequencies during each 10-min component). The first component of each daily session was considered to be a “warm up” component, and data were discarded. Data from the remaining pair of baseline components were averaged within each rat in a group and then across rats, and these data provided a measure of “Pre-Drug Baseline” (on Day 0) or “Daily-Baseline” (Days 1-7) ICSS performance. Similarly, test data from each pair of test components on Days 1-7 were averaged first within each rat and then across rats within each group to provide a measure of treatment effects on each test day. Daily baseline and test data from Days 1-7 were expressed as a percentage of the pre-drug baseline number of stimulations per component in each rat using the equation: % Pre-Drug Baseline Reinforcements per Component = (Daily-Baseline or Daily-Test Reinforcements per Component on a Test Day ÷ Pre-Drug Baseline Reinforcements per Component) × 100. Daily Baseline vs. Daily Test data for each group were analyzed using separate two-way ANOVAs, with treatment day as one factor and daily-baseline vs. daily-test performance as the second factor. A significant two-way ANOVA was followed by the Holm-Sidak post hoc test, and the criterion for significance in this and all other analyses was p<0.05.

A secondary and more granular measure of ICSS performance was the reinforcement rate in stimulations per frequency trial on Day 7, the final day of treatment in each group. Raw reinforcement rates for each rat from each trial were converted to “Percent Maximum Control Rate” (%MCR), with MCR defined as the mean of the maximal rates observed during any trial of the second and third components of the pre-drug baseline session (Day 0). Thus, %MCR values for each daily-baseline and test trial on Day 7 were calculated using the equation: %MCR = (reinforcement rate during a Day 7 baseline or test frequency trial ÷ MCR) × 100. %MCR values were then averaged across rats and analyzed by repeated-measures two-way ANOVA, with ICSS frequency as one factor and Day 7 baseline vs. test performance as the second factor. A significant two-way ANOVA was followed by the Holm-Sidak post hoc test.

Rats were weighed daily before treatment injections and behavioral sessions on Days 1-7, and body weight data for each rat on each day were expressed as a percent of the initial Pre-Drug weight on Day 1 using the equation % Pre-Drug Body Weight = (Weight on a Test Day ÷ Weight on Day 1) × 100. Data for each drug treatment were then analyzed by two-way ANOVA with treatment day as a within-subjects factor and daily treatment with IP acid or acid vehicle as a between-subjects factor. A significant ANOVA was followed by the Holm-Sidak post hoc test.

### IP Acid-Stimulated Stretching

#### Behavioral Procedure

To complement studies of IP acid-depressed ICSS, 5 rats (3 male, 2 female) were used in studies of IP 1.8% lactic acid-stimulated stretching as described previously (Negus et al. 2010; Rosenberg et al. 2013). Drugs were tested in the order ketorolac-nortriptyline-bupropion-citalopram-U69,593, and each drug was tested using a 7-day treatment protocol, with at least one week separating each test period. On Days 1 and 7 of each test period, the test drug was administered as a pretreatment to IP acid administration with the same doses and pretreatment times described above for ICSS studies. Immediately after the IP acid injection, rats were placed into an amber acrylic test chamber (31.0 × 20.1 × 20.0 cm) for a 30-minute observation period, and the number of stretches was recorded by an observer blind to the treatment. A stretch was operationally defined as an elongation of the subject’s body in a horizontal plane and contraction of abdominal muscles. On the intervening days of each test period (Days 2 - 6), rats were administered daily injections of the test drug at the same dose as on testing days; IP acid was not administered on Days 2-6. Baseline stretching elicited by IP acid in the absence of any test drug was also determined either one week prior to Day 1 or one week following Day 7 for each test period. Only rats with more than 5 stretches during all baseline assessments were included in the study. Nine rats failed to meet this criterion and were excluded from the study.

#### Data Analysis

The primary dependent variable was the number of stretches during each observation period in each rat. For each test period, baseline data were compared to Day 1 and Day 7 test data by one-way ANOVA. Baseline data across all five test periods were also compared by one-way ANOVA. A significant ANOVA was followed by Dunnett’s post hoc test.

### Drugs

Lactic acid, ketorolac HCl, nortriptyline HCl, and bupropion HCl were purchased from Sigma Chemical Company (St. Louis, MO). Citalopram HBr was purchased from Spectrum Chemical Manufacturing Corporation (Gardena, CA). U69,593 was obtained from the National Institute on Drug Abuse Drug Supply Program (Bethesda, MD). All drugs were dissolved in sterile water for IP injection.

## RESULTS

### Effects of control treatments ± IP acid on ICSS

For all rats in ICSS studies, the mean ± SEM pre-drug baseline number of reinforcements per component was 147.4±17.0, and the mean ± SEM pre-drug baseline maximum control rate was 59.7±7 reinforcements per trial. Figure 1 shows the time course of effects produced by repeated treatment with drug vehicle (panels a,d), the positive control ketorolac (panels b,e), or the negative control U69,593 (panels c,f) in the absence or presence of the IP acid noxious stimulus. Daily baseline ICSS performance was stable across days regardless of treatment (see open circles in all panels). Administration of drug vehicle with IP acid vehicle had no effect on ICSS (panel a), whereas administration of drug vehicle with IP acid produced repeatable ICSS depression on all seven days of the study (panel d; main effect of Baseline vs. Test; F (1,6)=47.47, p<0.001). A major goal of the study was to evaluate the effectiveness of the test drugs to block this sustained and repeatable IP acid-induced ICSS depression. The nonsteroidal anti-inflammatory drug ketorolac (10 mg/kg/day) administered with IP acid vehicle had no effect on ICSS (panel b), but consistent with its clinical effectiveness as an analgesic, ketorolac blocked IP acid-induced ICSS depression across all seven days of treatment (panel e). The kappa agonist U69,593 (0.18 mg/kg/day) also had no effect on ICSS when administered with IP acid vehicle (panel c), but U69,593 failed to block IP acid-induced ICSS depression on any day (panel f; main effect of Baseline vs. Test; F (1,5)=41.29, p=0.001). Figure 2 shows full ICSS frequency-rate curves before and after each treatment on Day 7. None of the treatments altered ICSS when administered before IP acid vehicle (panels a-c). IP acid administered after drug vehicle depressed ICSS by shifting the frequency-rate curve to the right (panel d, main effect of Baseline vs. Test; F (1,6)=16.75, p=0.006). Ketorolac blocked this effect (panel e), whereas U69,593 did not (panel f, main effect of Baseline vs. Test; F (1,5)=6.75, p=0.048).

**Figure 1:**
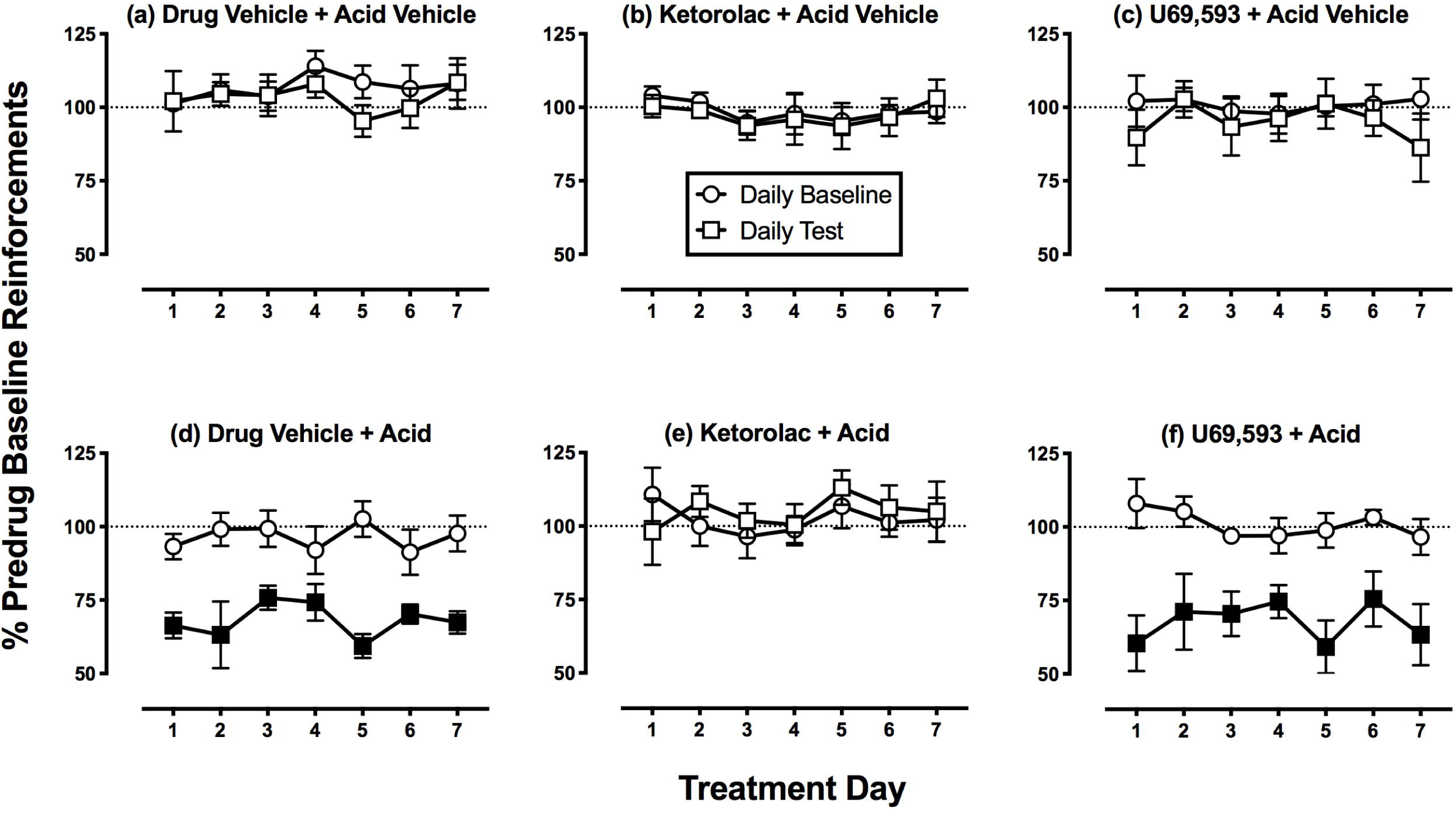
Time course of control-treatment effects on ICSS reinforcements per component. Horizontal axes: Time in days of treatment. Vertical axes: ICSS performance expressed as the % baseline number of pre-drug reinforcements earned per 10-min component. For all panels, circles indicate Daily Baseline data collected before the designated treatment, squares indicate Daily Test data collected after the designated treatment, and filled squares indicate significantly different from the Daily Baseline as indicated by two-way ANOVA followed by the Holm-Sidak post hoc test, p<0.05. All points show mean±SEM for N=6 rats (3 male and 3 female) except for the drug vehicle + acid group (N=7 rats, 3 male and 4 female).

**Figure 2:**
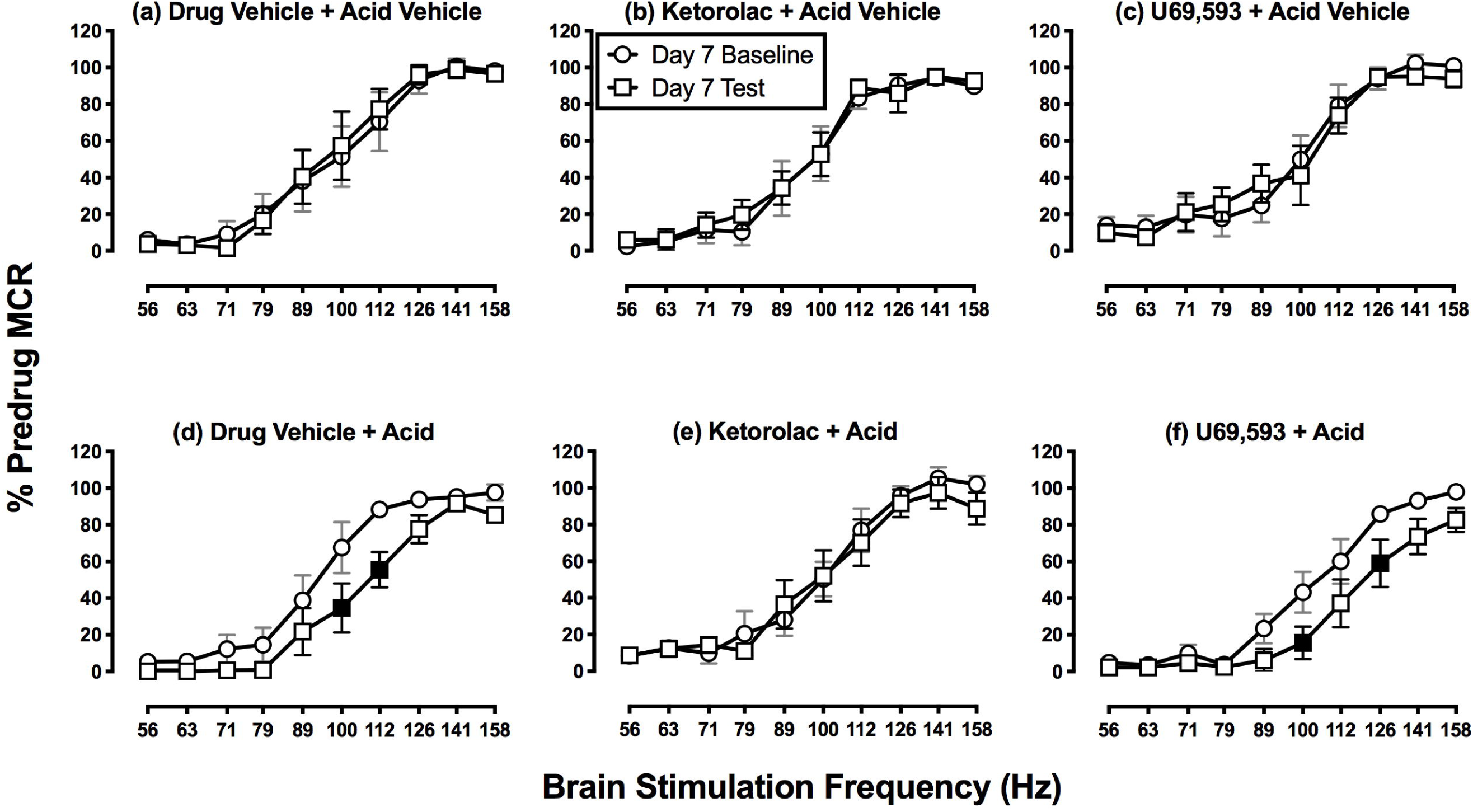
Control-treatment effects on Day 7 ICSS frequency-rate curves. Horizontal axes: frequency of brain stimulation in hertz (Hz; log scale). Vertical axes: ICSS performance expressed as the % pre-drug maximal control rate (MCR). For all panels, circles indicate Daily Baseline data collected before the designated treatment, squares indicate Daily Test data collected after the designated treatment, and filled squares indicate significantly different from the Day 7 Baseline as indicated by two-way ANOVA followed by the Holm-Sidak post hoc test, p<0.05. All points show mean±SEM for N=6 rats (3 male and 3 female) except for the drug vehicle + acid group (N=7 rats, 3 male and 4 female).

### Effects of antidepressants ± IP acid on ICSS

Figure 3 shows the time course of effects produced by repeated treatment with bupropion (panels a,d), nortriptyline (panels b,e), or citalopram (panels c,f) in the absence or presence of the IP acid noxious stimulus. Bupropion (3.2 mg/kg/day) produced modest but significant ICSS facilitation on Day 5 when it was administered with IP acid vehicle (panel a; main effect of Baseline vs. Test; F(1,5)=9.54, p=0.027), and it also blocked IP acid-induced ICSS depression on all treatment days (panel d). Nortriptyline (3.2 mg/kg/day) did not significantly alter ICSS when it was administered with IP acid vehicle (panel b), although there was a trend toward ICSS depression (main effect of Baseline vs. Test; F (1,6)=4.71, p=0.073). When nortriptyline was administered with IP acid, it produced a gradual onset of antinociceptive effect and fully blocked IP acid-induced ICSS depression by Day 7 (significant main effects of Treatment Day and Baseline vs. Test and a significant interaction; for interaction effect, F(6,36)=2.49, p=0.041). Citalopram (10 mg/kg/day) also had no effect on ICSS when administered with IP acid vehicle (panel c). When citalopram was administered with IP acid, it also produced a gradual onset of antinociceptive effect and fully blocked IP acid-induced ICSS depression by Day 7 (main effect of Baseline vs. Test; panel f, F(6,30)=3.52, p=0.009),. Figure 4 shows full ICSS frequency-rate curves before and after each treatment on Day 7. By this analysis, none of the treatments altered ICSS when administered before IP acid vehicle (panels a-c), and all of the treatments blocked acid-induced ICSS depression when drugs were administered before IP acid (panels d-f).

**Figure 3:**
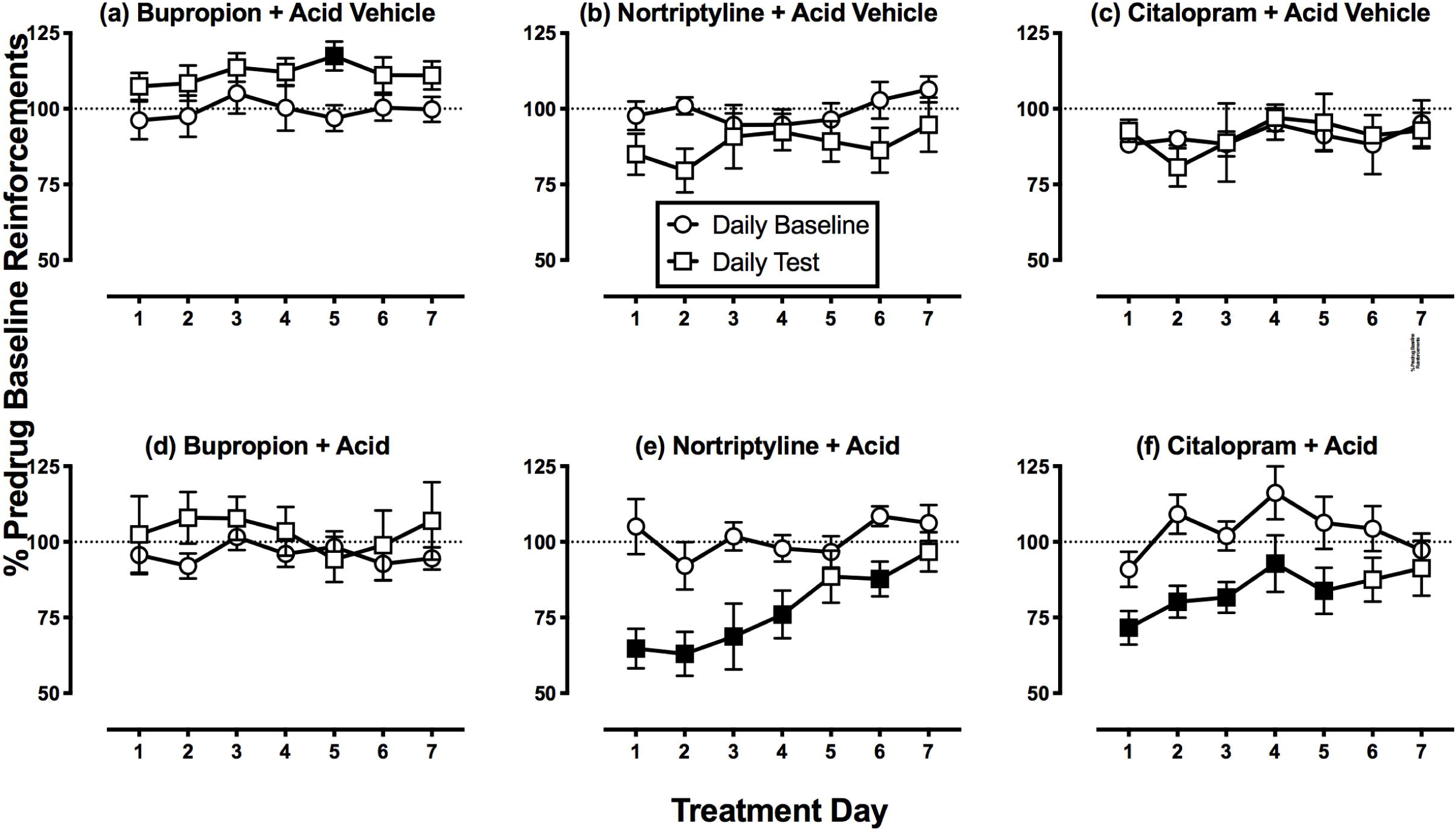
Time course of antidepressant effects on ICSS reinforcements per component. Horizontal axes: Time in days of treatment. Vertical axes: ICSS performance expressed as the % baseline number of pre-drug reinforcements earned per 10-min component. For all panels, circles indicate Daily Baseline data collected before the designated treatment, and squares indicate Daily Test data collected after the designated treatment. Filled squares indicate significantly different from the Daily Baseline as indicated by two-way ANOVA followed by the Holm-Sidak post hoc test, p<0.05. All points show mean±SEM for N=6 rats (3 male and 3 female) except for the nortriptyline + acid vehicle and nortriptyline + acid groups (N=7 rats, 4 male and 3 female).

**Figure 4:**
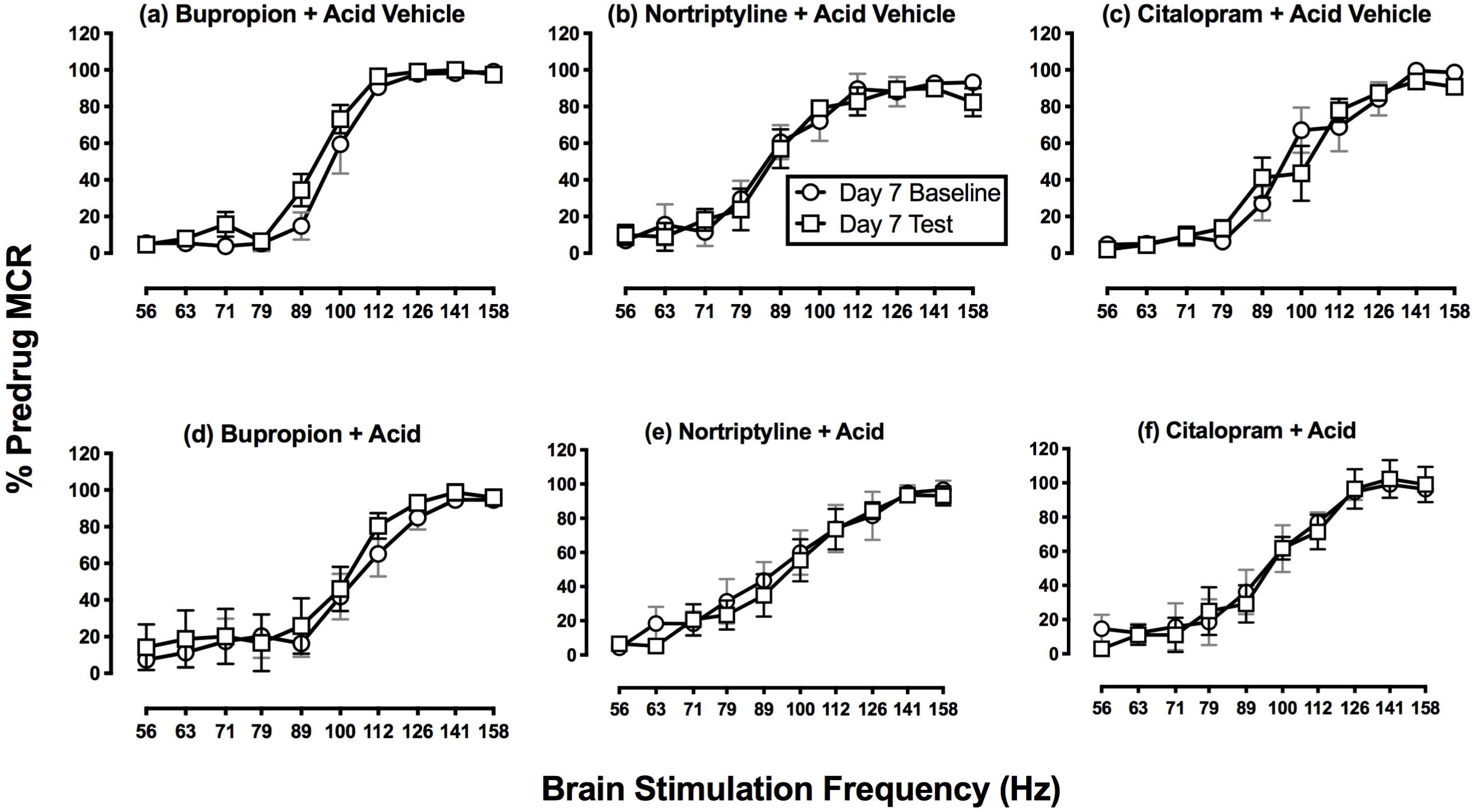
Antidepressant effects on Day 7 ICSS frequency-rate curves. Horizontal axes: frequency of brain stimulation in hertz (Hz; log scale). Vertical axes: ICSS performance expressed as the % pre-drug maximal control rate (MCR). For all panels, circles indicate Daily Baseline data collected before the designated treatment, squares indicate Daily Test data collected after the designated treatment, and filled squares indicate significantly different from the Day 7 Baseline as indicated by two-way ANOVA followed by the Holm-Sidak post hoc test, p<0.05. Two-way ANOVA identified no significant differences between Daily Baseline and Daily Test data for any treatment. All points show mean±SEM for N=6 rats (3 male and 3 female) except for the nortriptyline + acid vehicle and nortriptyline + acid groups (N=7 rats, 4 male and 3 female).

### Treatment effects on body weights

Figure 5 shows body weights during repeated administration of each treatment in the absence or presence of the IP acid noxious stimulus. Weight was relatively stable during repeated treatment with drug vehicle + IP acid vehicle, and repeated treatment with drug vehicle + IP acid caused a gradual loss of body weight (panel a, significant main effects of Treatment Day and Acid Treatment and a significant interaction; for interaction effect F(6,66)=8.36, p<0.001). Accordingly, the drugs could be evaluated for their effectiveness to block IP acid-induced weight loss. Ketorolac (panel b), bupropion (panel d), and citalopram (panel f) all blocked IP acid-induced weight loss. Conversely, U69,593 (panel c) and nortriptyline (panel e) produced weight loss whether they were administered in the absence or presence of IP acid (main effect of Treatment Day was F(6,60)=9.93, p<0.001, for U69,593, and F(6,72)=10.95, p<0.001, for nortriptyline). Thus, U69,593 and nortriptyline failed to block IP acid-induced weight loss.

**Figure 5:**
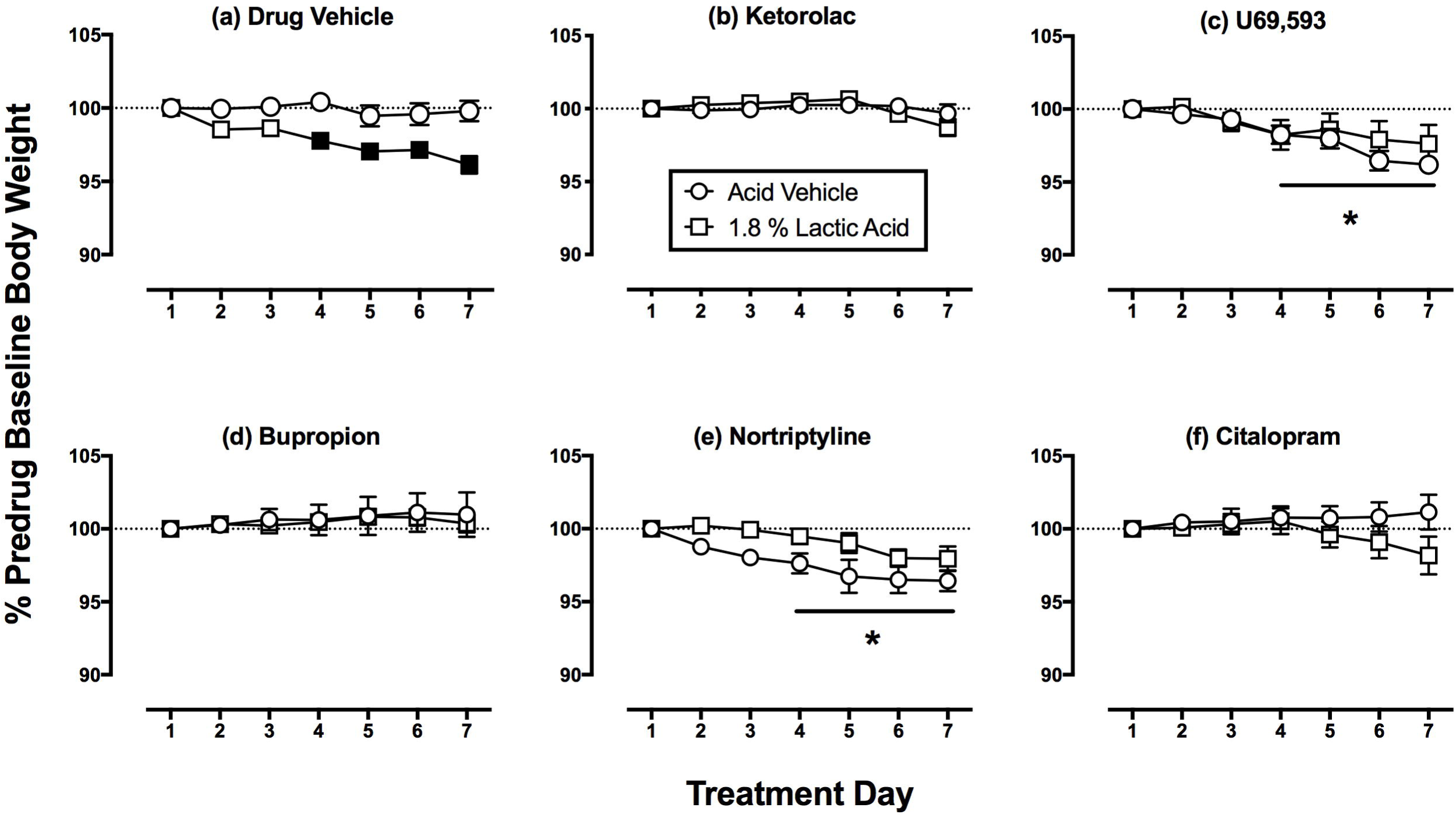
Time course of treatment effects on body weights. Horizontal axes: Time in days of treatment. Vertical axes: Body weight expressed % pre-drug baseline body weight. For all panels, circles indicate effects of each treatment administered with acid vehicle, and squares indicate effects of each treatment administered with acid. Filled squares indicate significantly different from the Acid-Vehicle group as indicated by a significant interaction between Treatment Day and Acid Treatment, whereas asterisks indicate significantly different from Day 1 due to a significant main effect of Treatment Day (two-way ANOVA followed by the Holm-Sidak post hoc test, p<0.05). All points show mean±SEM for N=6 rats (3 male and 3 female) except for the nortriptyline + acid vehicle and nortriptyline + acid groups (N=7 rats, 4 male and 3 female).

### Treatment effects on IP acid-stimulated stretching

Figure 6 shows the effects of repeated treatment with each drug on IP acid-stimulated stretching behavior. There was no difference in baseline stretching across test periods. Only ketorolac [F(1.23,4.90)=31.58; p=0.002] and U69,593 [F(1.22,4.88)=37.14; p=0.002] significantly decreased stretching, and both drugs significantly reduced stretching on both Day 1 and Day 7 of repeated treatment.

**Figure 6:**
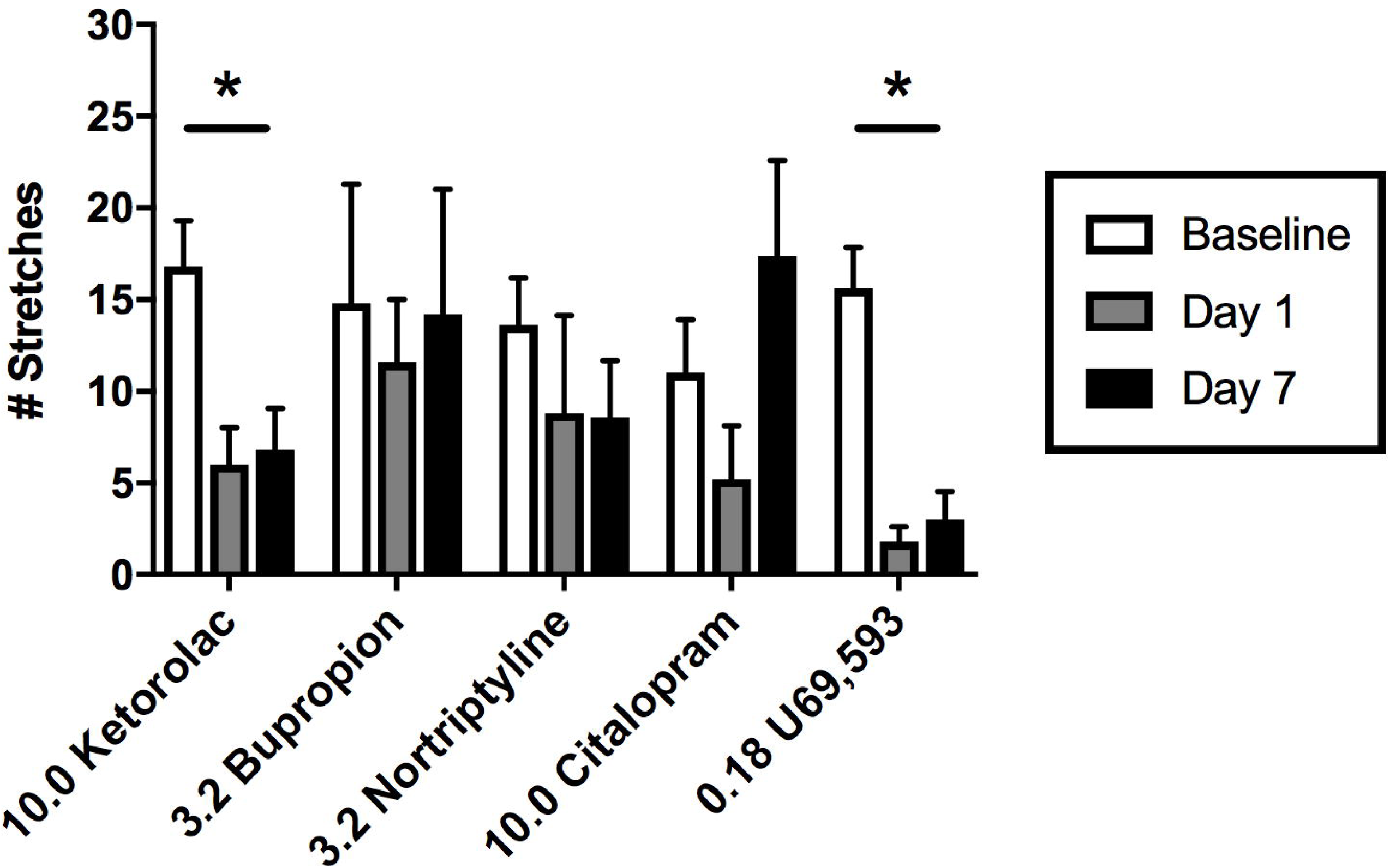
Effects of test compounds on acid-stimulated stretching. Horizontal axis: drug and dose in mg/kg/day. Vertical axis: Number of stretches in 30-min test session. Each bar displays mean±SEM for N=5 rats (3 male, 2 female). Asterisk and line denote statistical significance between baseline and test Days 1 and 7 by one-way ANOVA followed by a Dunnett’s post hoc (P<0.05). Drugs are listed left to right in the order in which they were tested.

## DISCUSSION

This study evaluated effects of monoamine transporter inhibitors on pain-related behavioral depression produced in male and female rats by IP acid as a visceral noxious stimulus. Repeated daily administration of IP acid for seven consecutive days decreased both body weight and positively reinforced operant responding in an intracranial self-stimulation (ICSS) procedure. This acid-induced depression of ICSS and body weight provided an opportunity to compare antinociceptive effectiveness of repeated test-drug administration.

Results are summarized in Table 1, and there were three main findings. First, consistent with the clinical effectiveness of NSAIDs for treatment of inflammatory pain, ketorolac blocked acid-induced depression of both ICSS and body weight. Second, consistent with the poor analgesic effectiveness of centrally acting kappa opioid receptor agonists, U69,593 failed to block these acid effects. Lastly, the monoamine-transporter-inhibitor antidepressants bupropion, nortriptyline, and citalopram all alleviated acid effects, though to different degrees and with different time courses. The DAT/NET-selective inhibitor bupropion produced a ketoprofen-like blockade of acid-induced ICSS depression and weight loss throughout the seven days of treatment. By contrast, the NET-selective inhibitor nortriptyline and SERT-selective inhibitor citalopram produced more gradual blockade of acid-induced ICSS depression that emerged only after repeated treatments, and nortriptyline failed to block acid-induced weight loss.

**TABLE 1:**
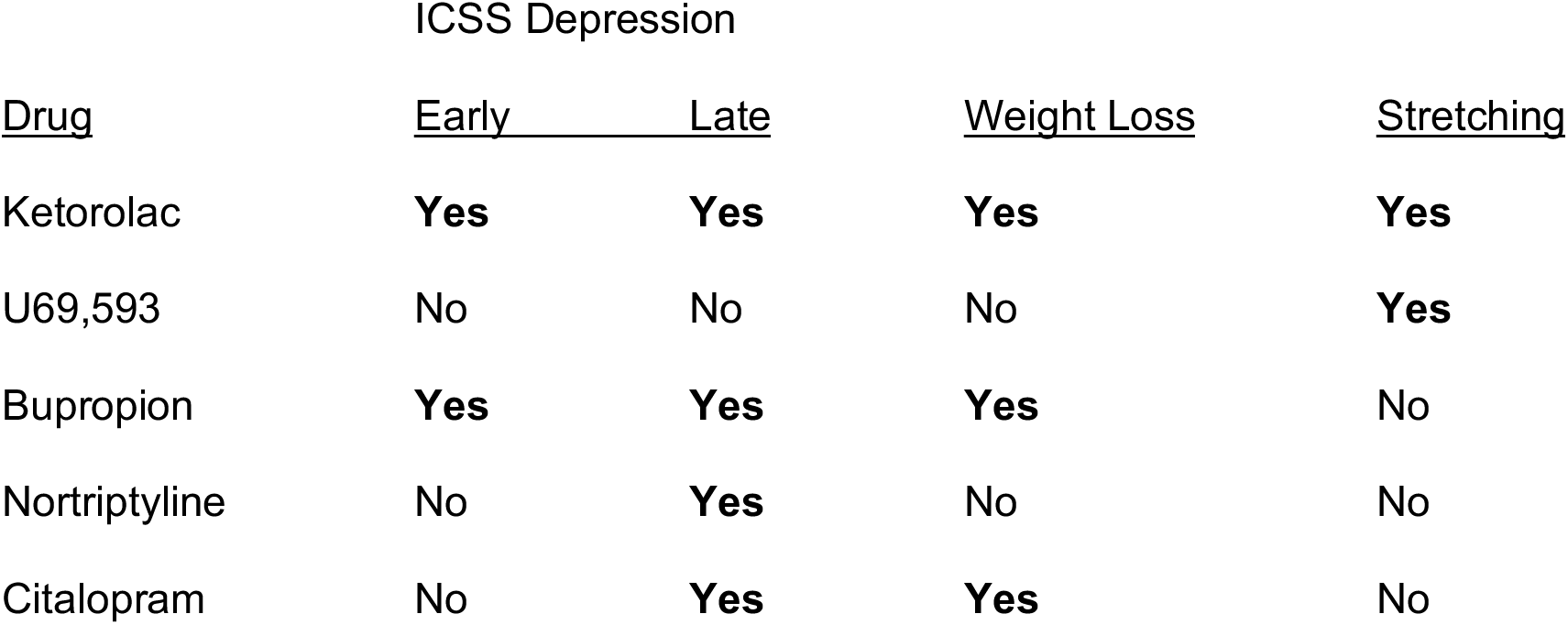
Summary of drug effects on acid-induced depression of ICSS and body weight and acid-induced stimulation of stretching.

### Effects of repeated IP acid

ICSS is one type of positively reinforced operant behavior that can be reduced by some noxious stimuli that are thought to produce pain states in rodents (Negus 2013). We and others have shown that acute IP acid can serve as a visceral noxious stimulus sufficient to produce ICSS depression as a model of pain-related behavioral depression in rats (Brust et al. 2016; Pereira Do Carmo et al. 2009). The present results confirm and extend our previous report that repeated daily administration of IP acid can also produce repeatable ICSS depression in male rats and extends this finding to female rats (Miller et al. 2015). Although IP acid effects on food consumption were not evaluated directly in this study, prior studies have shown that IP acid can decrease feeding in both rats and mice (Kwilasz and Negus 2012; Stevenson et al. 2006), and consistent with these findings, the present study found that repeated IP acid resulted in weight loss. This effectiveness of repeated IP acid to produce repeated ICSS depression and weight loss provides an opportunity to evaluate effects of repeated treatment with candidate analgesic drugs. We reported previously that the mu opioid receptor agonist morphine retained its antinociceptive effectiveness to alleviate IP acid-induced ICSS depression during 7 days of repeated morphine treatment (Miller et al. 2015). The present study used this same procedure to evaluate effects of repeated treatment with monoamine-transporter-inhibitor antidepressants in comparison to the NSAID ketorolac and the kappa opioid receptor agonist U69,593 as positive and negative controls, respectively.

### Effects of ketorolac and U69,593

Ketorolac was effective to alleviate both IP acid-induced stimulation of stretching and depression of ICSS on the first day of ketorolac administration, and it retained its effectiveness on both endpoints throughout 7 days of chronic treatment. Ketorolac also blocked IP acid-induced weight loss. These findings are consistent with the analgesic effectiveness of ketorolac in human and veterinary medicine (Litvak and McEvoy 1990; Matava 2018; Mathews et al. 1996) and also agree with our previous report that acute ketorolac treatment blocked acute IP acid-induced ICSS depression (Moerke et al. 2019). Ketorolac and other NSAIDs can produce gastric ulceration upon repeated exposure, and this in turn has the potential to reduce body weight; however, the dosing regimen used here in the absence or presence of repeated IP acid was not sufficient to alter body weight. U69,593 also blocked IP acid-induced stimulation of stretching both on the first day of its administration and after repeated treatment for 7 days; however, U69593 failed to block IP acid-induced ICSS depression at any time, and it also failed to block weight loss. These results are consistent with other evidence to suggest that acute treatment with U69,593 and other centrally acting kappa opioid receptor agonists reduce pain-stimulated behaviors but fail alleviate pain-related behavioral depression (Bagdas et al. 2016; Brust et al. 2016; Lazenka et al. 2018; Negus et al. 2010; Negus et al. 2015; Negus et al. 2012). The present results extend these results to suggest that antinociceptive effects do not emerge with repeated treatment. Taken together, these findings are consistent with clinical evidence to suggest that centrally acting kappa opioid receptor agonists are not effective analgesics in humans (Lazenka et al. 2018; Pande et al. 1996).

### Effects of bupropion, nortriptyline, and citalopram

Like ketoprofen, bupropion blocked IP acid-induced ICSS depression and weight loss throughout the 7 days of treatment. The effects of bupropion on Day 1 replicate our earlier report that acute bupropion treatment alleviates acute IP acid-induced ICSS depression in male rats (Rosenberg et al. 2013), and the sustained effectiveness of bupropion in the present assay of repeated IP acid-induced ICSS depression agrees with the sustained effectiveness of repeated 7-day bupropion to alleviate formalin-induced ICSS depression (Leitl and Negus 2016). The bupropion dose tested in this study (3.2 mg/kg) did not reduce IP acid-stimulated stretching after either acute or repeated treatment; however, we reported previously that higher bupropion doses (10-32 mg/kg) do produce a dose-dependent decrease in stretching. Taken together, these findings provide preclinical evidence for analgesic effectiveness of bupropion and suggest that it may be especially effective to alleviate behavioral depressant signs of pain. Additionally, these preclinical findings agree with evidence for clinical analgesic effectiveness of bupropion and the mechanistically similar mixed DAT/NET inhibitor methylphenidate under at least some conditions (Pud et al. 2017; Shah and Moradimehr 2010).

When bupropion was administered alone in the absence of the IP acid noxious stimulus, it weakly but significantly increased ICSS. This is consistent with our previous findings with this relatively low dose of 3.2 mg/kg bupropion (Leitl and Negus 2016; Rosenberg et al. 2013), and higher bupropion doses produce much larger magnitudes of ICSS facilitation similar to those produced by abused monoamine transporter inhibitors such as cocaine, methylphenidate, and methylenedioxypyrovalerone (MDPV) (Bonano et al. 2014; Kornetsky and Esposito 1979; Lazenka and Negus 2017). ICSS facilitation is suggestive of abuse potential (Carlezon and Chartoff 2007; Negus and Miller 2014; Wise 1996), and although bupropion is not currently scheduled by the Drug Enforcement Administration in the United States, it does produce abuse-related effects in other preclinical procedures in laboratory animals (e.g. drug self-administration (Lamb and Griffiths 1990; Nicholson et al. 2009)), and abuse has been observed clinically in the United States and elsewhere (Dagan and Yager 2018; Stall et al. 2014; Stassinos and Klein-Schwartz 2016). Thus, while results of the present study are consistent with other evidence to suggest that bupropion has potential as a non-opioid analgesic, its potential for abuse should also be considered.

In contrast to bupropion, neither nortriptyline nor citalopram alleviated IP acid-induced ICSS depression on the first day of treatment. This agrees with our previous finding that acute treatment with these compounds, and with the related compounds nisoxetine (a NET-selective inhibitor) and clomipramine (a SERT-selective inhibitor), failed to block acute IP acid-induced ICSS depression (Rosenberg et al. 2013). However, after 7 days of repeated treatment, both compounds fully blocked IP acid-induced ICSS depression. This emergence of antinociceptive effectiveness is consistent with clinical evidence that repeated treatment with NET- and/or DAT-inhibitor antidepressants is usually required before analgesic effectiveness is apparent (Sutherland et al. 2018). Moreover, the emergence of antinociceptive effectiveness during repeated treatment with nortriptyline or citalopram + IP acid was not accompanied by the emergence of ICSS facilitation in rats treated with nortriptyline or citalopram + acid vehicle. The failure of these compounds to facilitate ICSS even after repeated treatment is consistent with their low abuse liability (Negus and Miller 2014), and also suggests that blockade of IP acid-induced ICSS depression cannot be attributed to a nonselective increase in ICSS responding. Rather, these results suggest that repeated treatment with nortriptyline and citalopram engaged processes that selectively attenuated pain-related ICSS depression.

Repeated citalopram also significantly attenuated IP acid-induced weight loss, suggesting that citalopram alleviated pain-related depression of feeding as well as ICSS. Nortriptyline effects on IP acid-induced feeding were more difficult to interpret given weight loss after nortriptyline alone; however, nortriptyline did not exacerbate IP acid-induced weight loss. Additionally, although neither compound significantly reduced IP acid-stimulated stretching at the doses used in the the present study, we reported previously that both compounds decreased IP acid-stimulated stretching at higher doses, and other studies have also reported antinociceptive of these and related compounds in assays of pain-stimulated behaviors elicited by chemical noxious stimuli (Aoki et al. 2006; Ardid et al. 1992; Leventhal et al. 2007; Rojas-Corrales et al. 2003).

## Conclusions

These results support the utility of antidepressant drugs to treat pain-related behavioral depression and suggest that the DAT/NET inhibitor bupropion might have a faster onset and higher effectiveness than NET- or SERT-selective inhibitors. Mechanisms that underlie emergent antinociception with NET and SERT inhibitors require further study.

## Acknowledgements

This work was supported by grants R01NS070715 (SSN) and F30CA213956 (LPL) from the National Institutes of Health. The authors have no conflicts of interest to report.

